# Dominant Role of Stochastic Processes in Soil Fungal Communities in Pioneer Forests at a Regional Scale

**DOI:** 10.1101/2023.02.27.530225

**Authors:** Xiaowu Man, Qingchao Zeng, Meng Zhou, Francis M. Martin, Feng Zhang, Honggao Liu, Yuan Yuan, Yucheng Dai

## Abstract

Soil fungal community assembly is driven by deterministic and stochastic processes. However, the contribution of these mechanisms to structure the composition of fungal communities of forest soils at the regional scale is poorly known. Here, we investigate the relative importance of deterministic and stochastic processes on fungal community composition by rDNA ITS metabarcoding in a *Populus davidiana* pioneer forests along spatial-temporal gradients. We also assessed the impact of elevation and seasonality. The soil fungal richness of *P. davidiana* pioneer forests was significantly affected by elevation and less affected by season. Similarly, the variation in the fungal community composition according to the elevation was greater than the effect of seasonality. The fungal community composition showed a significant distance-decay relationship. Variation partitioning analysis showed that plant variables explained the soil fungal community variation. Through null model analysis, we found that stochastic processes were dominant in the soil fungal community assembly. However, the relative importance of ecological processes, including dispersal, selection, and drift, was not consistent across the four soil fungal community assemblies. In addition, the undominated fraction (including weak selection, weak dispersal, diversification and drift) had a high relative contribution to the soil fungal community assembly process in the *P. davidiana* pioneer forest. In summary, our results demonstrated that plant variables and the undominated fraction dominate the deterministic and stochastic processes driving soil fungal community assemblies in a *P. davidiana* pioneer forest at the regional scale, which provides new perspectives for the regional scale studies of soil fungi.

**IMPORTANCE:** Elevation and seasonality are important factors driving the composition of soil microbiota. Due to the tight interactions of soil fungi with their host trees in forest ecosystems, the spatial variation of soil fungal community is often linked to the variation in the composition of dominant tree species. We compared the responses of soil fungal communities to seasonal and spatial changes at four levels in a temperate poplar forest dominated by a single tree species under elevation changes. Elevation had a higher impact than seasonality on the soil fungal beta diversity. Even when the shift in dominant tree species was limited, vegetation factors still impact soil fungal community variations. The dominant role of homogeneous selection and drift in fungal community assemblies, except for ectomycorrhizal fungi, was further discovered.

## INTRODUCTION

The forest ecosystem is one of the most important terrestrial ecosystems, providing key environmental contributions for the biosphere, such as being a carbon sink, protecting biodiversity, protecting the soil, and providing wood materials and resources (1). As an indispensable part of forest ecosystems, soil microorganisms usually play major ecological roles. In particular, soil fungal communities affect the forest ecosystem processes by participating in organic matter decomposition, mutualistic symbiosis, or plant diseases. These fungal communities respond to changes in biotic and abiotic factors (2). Specifically, trees, as the dominant plants in forest ecosystems, affect the soil fungi community by changing soil coverage and structure, regulating soil temperature and humidity, and affecting understory productivity (3). In addition, the tree root structure and secretions directly affect the soil fungal community composition, especially mycorrhizal fungal communities, by changing soil properties and selecting host-specific symbiotic fungi (4, 5, 6). The impact of abiotic factors, which include soil, climate, and other environmental variables, on soil microorganisms also modulate the richness and composition of fungal communities (7). The effects of soil pH, carbon-nitrogen ratio, and phosphorus content on the soil fungal community composition have been well documented (8, 9, 10, 11, 12, 13, 14).Climatic factors could also significantly regulate the soil fungal community. Precipitation can directly affect the growth of soil fungi by changing soil moisture (15, 16, 17). At the same time, runoff caused by precipitation causes the distribution of nutrients and changes in root biomass (18, 19, 20), indirectly affecting soil fungal communities’ occurrence and spread (21, 22, 23, 24, 25). Similarly, temperature changes can also affect the input of organic matter by changing plant communities and productivity (26), thereby indirectly affecting the occurrence of soil fungal communities (27, 28).

Although the dynamic changes of soil fungal communities are impacted by biotic and abiotic factors, they can have variable effects across different ecosystems, on different fungal communities, and at different scales. For example, it has been reported that climate factors, followed by soil factors and spatial patterns, are the best predictors of total soil fungal richness and community composition on a global scale. The pH, distance from the equator, and host richness were strong predictors of ectomycorrhizal fungal richness (29, 30). At the continental scale, soil fungal community variations are mainly affected by environmental variables, such as pH and precipitation (31). At the regional scale, studying soil fungal communities and interpreting their dynamics are more complex and diverse. Among them, there are only a few reports addressing the spatial-temporal variation in soil fungal communities along elevation gradients and between seasons. To our best of our knowledge, these studies did not provide a unified explanation for the spatial and temporal variation of soil fungal communities across different elevations and seasons. For example, clear elevation patterns, in which soil fungal richness decreased with elevation, were observed in certain studies (32) but not in others (33). Additional work has shown that the vertical distribution patterns of soil fungal communities observed in these studies were usually explained by environmental factors that change along elevation gradients. These include changes in soil moisture and organic carbon at different elevations, resulting in soil fungal diversity and community composition patterns changing with elevation (34, 35). The seasonal patterns of soil fungal communities also appear to be inconsistent, according to recent reports. In some studies, the soil fungal communities showed significant seasonal variation (36, 37). Others, in contrast, have reported that the season did not affect the variation of the soil fungal community (38, 39, 40). Moreover, seasonal patterns are usually explained by seasonal variations in precipitation, and temperature, among others (41, 42, 43). Moreover, the impact of the host on soil fungal communities cannot be ignored (4, 44, 45, 46, 47). A number of studies have shown that at the regional scale, along the elevation gradient, changes in host species and richness will lead to soil fungal communities’ elevation pattern (48). Therefore, the soil fungal communities’ elevation and seasonal variation, as affected by the host species, is still a proposition worthy of further study.

In addition to deterministic processes controlling microbial community structure proposed by traditional niche-based theoretical assumptions, stochastic processes proposed by the neutral theory of evolution have been widely discussed in microbial ecology recently to control microbial community structure (49, 50). It is generally believed that stochastic processes affect the assembly of soil microbial communities on a large scale (51). At the regional scale, deterministic processes usually play a decisive role in the assembly of soil fungal communities (32). However, recent studies have shown that stochastic processes are also important in soil microbial community assembly at the regional scale (52, 53, 54). Furthermore, conclusions differ on the relative importance of diffusion limitation, selective diffusion, homogeneous selection, heterogeneous selection, and undominated fraction (including weak selection, weak dispersal, diversification and drift) in the soil fungal community assembly process, especially with regard to different functional fungal community assemblies (55, 51). Therefore, exploring which process dominates the assembly of soil fungal communities with different functions at the regional scale can help us better understand the underlying governing mechanisms.

As the mechanisms underlying dynamic changes of different soil functional fungal communities between elevations and seasons have not yet been fully elucidated, and the ecological process of community assembly remains controversial, we designed this study to investigate the dynamics of fungal communities, such as mycorrhizal and saprotrophic species, at different elevations and seasons. In addition, due to the host tree’s influence on soil fungal communities (56, 57), we explored the dynamics of forest soil fungal communities associated to a single host tree species, *P. davidiana*.

*P. davidiana* is widely distributed inin temperate forests in China. It is a deciduous tree species and an important source of timber. It provides excellent materials, has a straight trunk, and excellent physiological characteristics such as cold tolerance, drought resistance, and barren tolerance. It can adapt to different climates and environments, and it can grow on slopes, ridges, and valleys. Its wide adaptation potential is in part due to its ability to form symbiotic mycorrhizas with soil fungi, which can help them adapt under stress conditions (58, 59, 60), so they are usually colonized in forests as pioneer species. As a dual-mycorrhizal tree species, the roots of *P. davidiana* can form associations with both ectomycorrhizal and endomycorrhizal fungi. However, However, the information available on the soil fungal community associated to this tree is limited (61). It is unclear how soil fungal communities, especially ectomycorrhizal and endomycorrhizal fungal communities, are distributed and how they respond to environmental changes.

Therefore, in this study, we comprehensively investigated the soil fungal community of *P. davidiana* in Xinglong Mountain. *P. davidiana* is colonized as a pioneer tree species in the high-elevation mountain forest of the Xinglong Mountain area of Lanzhou City, Gansu Province, northwest China. To explore the soil fungi and mycorrhizal fungi diversity, community composition, dynamic changes, and community assembly process in different *P. davidiana* forests in this area, we used high-throughput sequencing approaches. We hypothesized that: (i) the soil fungal richness and community composition of poplar forests at different elevations would vary due to the environmental spatial and temporal heterogeneity; (ii)the responses of total soil fungi and different types of mycorrhizal fungi to spatiotemporal variations in richness and community composition would be variable; (iii) The assembly processes of soil fungi and different types of mycorrhizal fungal communities would be different. Understanding these can help us better understand the reasons for *P. davidiana* wide adaptation in various environments and can also help us understand the mechanisms governing microbial community composition.

## RESULTS

### Site information and variation of environmental variables between elevations and seasons

The site information and landscape information of low elevation (XL2300), middle elevation (XL2500) and high elevation (XL2600) were collected (Figure 1). The variation of soil physical and chemical properties among elevation was significant (available phosphorus (AP): *R^2^* = 0.582, *P* <0.01; cation exchange capacity (CEC): *R^2^* = 0.151, *P* < 0.01; organic carbon (OC): *R^2^* = 0.513, *P* < 0.01; pH: *R^2^* = 0.596, *P* < 0.01). Soil AP and CEC were significantly higher in XL2500 and XL2600 compared to XL2300. Soil OC in XL2300 was significantly higher than that in XL2500 and XL2600. The highest soil pH was observed at XL2500. However, the soil in all three elevations was weakly alkaline (Figure 2). Among the 4 environmental variables describing above-ground vegetation condition, diameter at breast height of the tree (tree DBH)(*R^2^* =0.857, *P* < 0.01) and ground primary productivity (GPP)(*R^2^* = 0.887, *P* < 0.01) differed significantly between zones, with tree DBH decreasing significantly from low to high elevations; GPP was highest at XL2300 and lowest at XL2500. enhanced vegetation index (EVI)(*R^2^* = 0.764, *P* < 0.01) and gap-filled of ground primary productivity (GPP GF)(*R^2^* = 0.60, *P* < 0.01) were significantly affected by the different seasons and not affected by elevation zones. Except for GPP GF in XL2600, EVI and GPP GF in other zones were significantly higher during the summer than in autumn (Figure 2). The GPP at XL2300 also showed significant seasonal differences (higher in summer than in autumn). In contrast, GPP was not significantly affected by the season in the other two zones (Figure 2).

**FIG 1.**
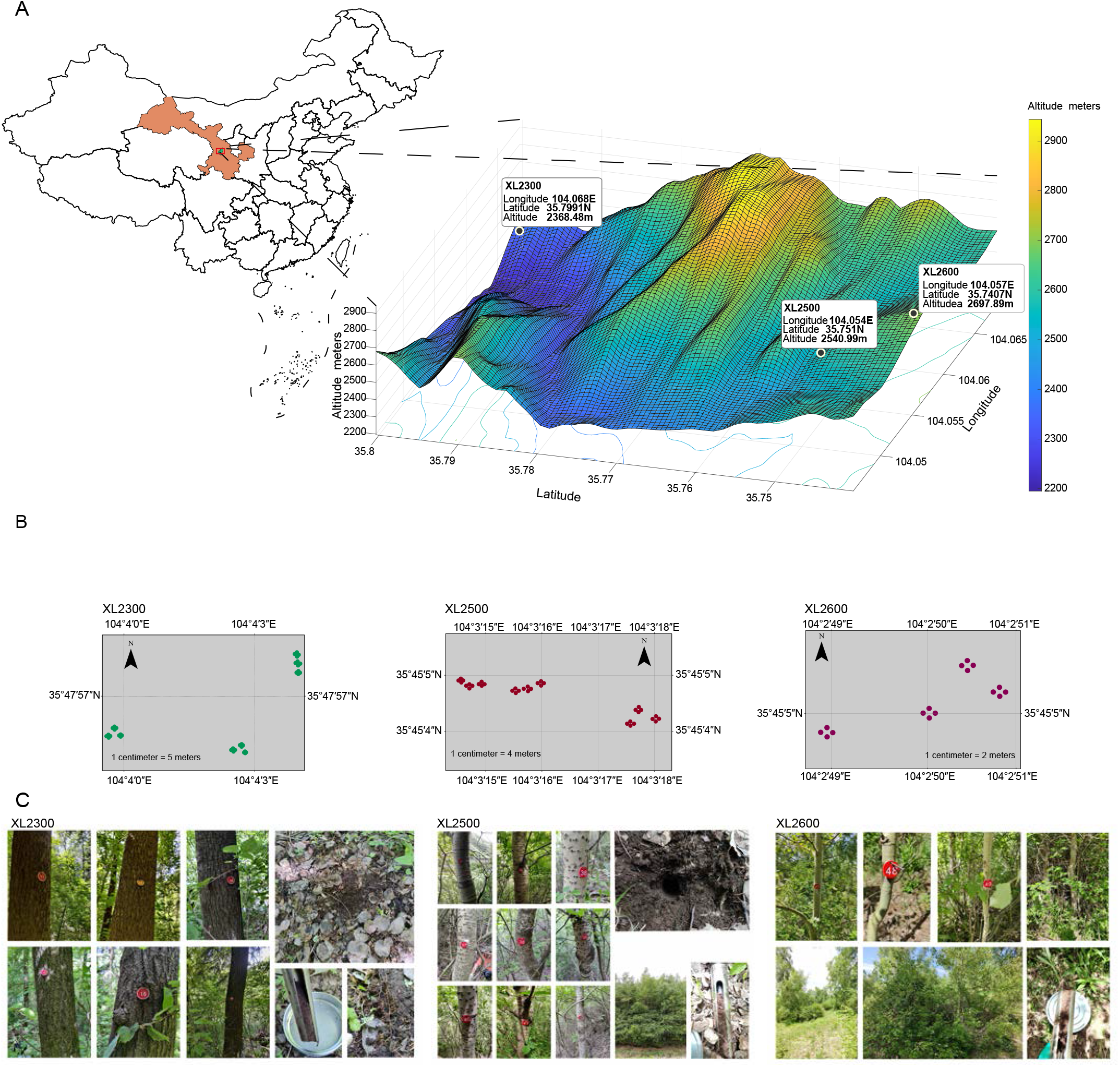
Distribution of transects at different elevations and distribution of sampling points in Xinglong Mountain. (A) The study area location and the distribution of each elevation. (B) Detailed sampling point distribution map from each elevation. (C) Landscape photos and sampling detail photos from each elevation. The 3D map was made by Matlab, and the detailed sampling point map of each elevation transect was made by ArcMap. The map of China was obtained from https://geo.datav.aliyun.com/areas_v2/bound/100000_full.json and was visualized by the R package sf.

**FIG 2.**
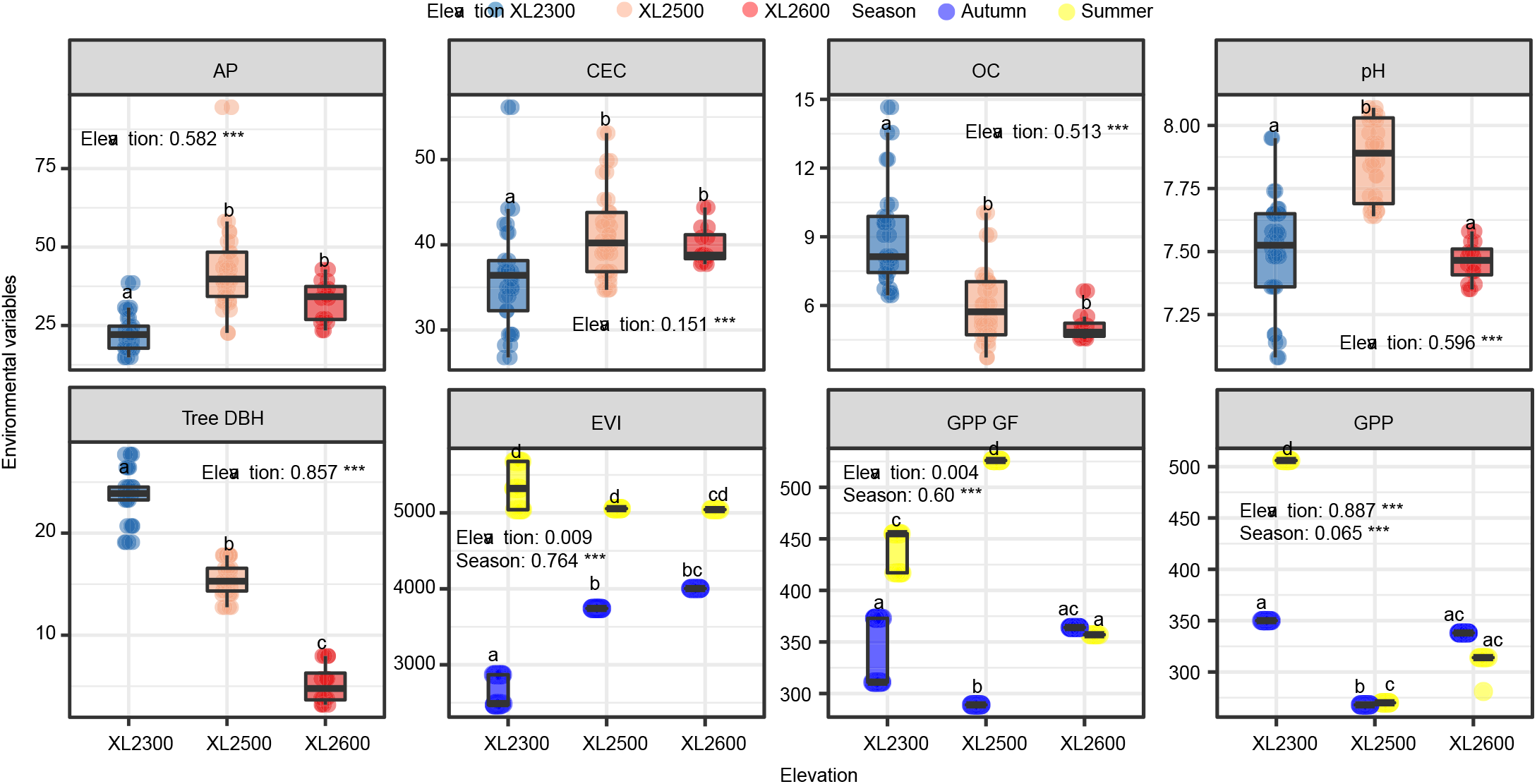
Distribution of environmental variables between seasons and zones. A nonparametric test (Scheirer-Ray-Hare test) was performed in the two-way factorial design and indicated significant differences in EVI, GPP, and GPP GF between elevations and seasons. The Kruskal-Wallis rank sum test indicated significant differences in four soil variables and tree DBH between zones. The number indicated after the elevation and season correspond to *R^2^*, representing the variation in environmental variables explained by season and zone, ***, *P* < 0.001; AP, soil available phosphorus; CEC, Soil cation exchange capacity; OC, soil organic carbon; pH, soil acidity and alkalinity, Tree DBH, DBH of the sampled tree; EVI, enhanced vegetation index; GPP, ground primary productivity; GPP GF, gap-filled of ground primary productivity.

### Spatiotemporal distribution of soil fungal community

For the Illumina NovaSeq sequencing, 5,790,342,10,886,725, and 13,458,869 high-quality sequences with 1,710, 3,699, and 1,742 operational taxonomic units (OTUs) for the ITS1, ITS2, and AMF regions were obtained, respectively. A total of 30,135,936 high-quality sequences corresponding to 7,151 OTUs were obtained from the three fungal regions. Through classification, soil fungi were mainly attributed to Agaricomycetes, Pezizomycetes, Ar-chaeorhizomycetes, Leotiomycetes, Glomeromycetes and Mortierellomycetes at the class level (Fig. S2A,B). The relative abundances of Agaricomycetes, Pezizomycetes and Glomeromycetes were significantly different among elevations (Fig. S2C). The relative abundance of Pezizomycetes, Archaeorhizomycetes, Leotiomycetes, Glomeromycetes and Mortierellomycetes varied significantly among seasons (Fig. S2D). At the OTU level, XL2300 has the most specific OTUs (Fig. S2E), and XL2600 has the least specific OTUs.There were 1172 and 1121 specific OTUs in autumn and summer, respectively (Fig. S2F). However, the relative abundance of specific OTUs relative to different elevations and seasons were low (Fig. S2E,F). After functional prediction, there were 1,377 OTUs classified as total mycorrhizal fungi, 729 OTUs classified as ectomycorrhizal fungi, and 648 OTUs classified as endomycorrhizal fungi. The soil fungal alpha diversity variation was mainly influenced by forest elevation (Figure 3). In particular, the richness of soil fungi varied significantly between different elevation (total fungi: *R^2^* = 0.090, *P* < 0.001; total mycorrhizal fungi: *R^2^* = 0.161, *P* < 0.001; ectomycorrhizal fungi: *R^2^* = 0.057, *P* < 0.01; endomycorrhizal funi: *R^2^* = 0.415, *P* < 0.001). The richness of endomycorrhizal fungi increased significantly with elevation (Figure 3, Fig. S3). Similarly, the richness of total fungi and total mycorrhizal fungi also exhibited an elevation pattern with increasing elevation in autumn; however, no obvious elevation pattern was observed in the summer (Figure 3, Fig. S3). The richness of ectomycorrhizal fungi was higher in XL2300 and XL2500 than in XL2600, especially during the summer. The Shannon diversity of total soil fungi and endomycorrhizal fungi varied significantly among elevational zones, showing an elevational pattern that increased significantly with elevation (Figure 3, Fig. S3). The effect of season on the soil fungi alpha diversity was limited. Only the richness (total fungal:*R^2^* = 0.144, *P* < 0.001) and the Shannon diversity (total fungal: *R^2^* = 0.062, *P* < 0.001; ectomycorrhizal fungal: *R^2^* = 0.045, *P* < 0.01) of the total and ectomycorrhizal fungi were significantly different between seasons, being significantly higher in autumn than in summer (Figure 3).

**FIG 3.**
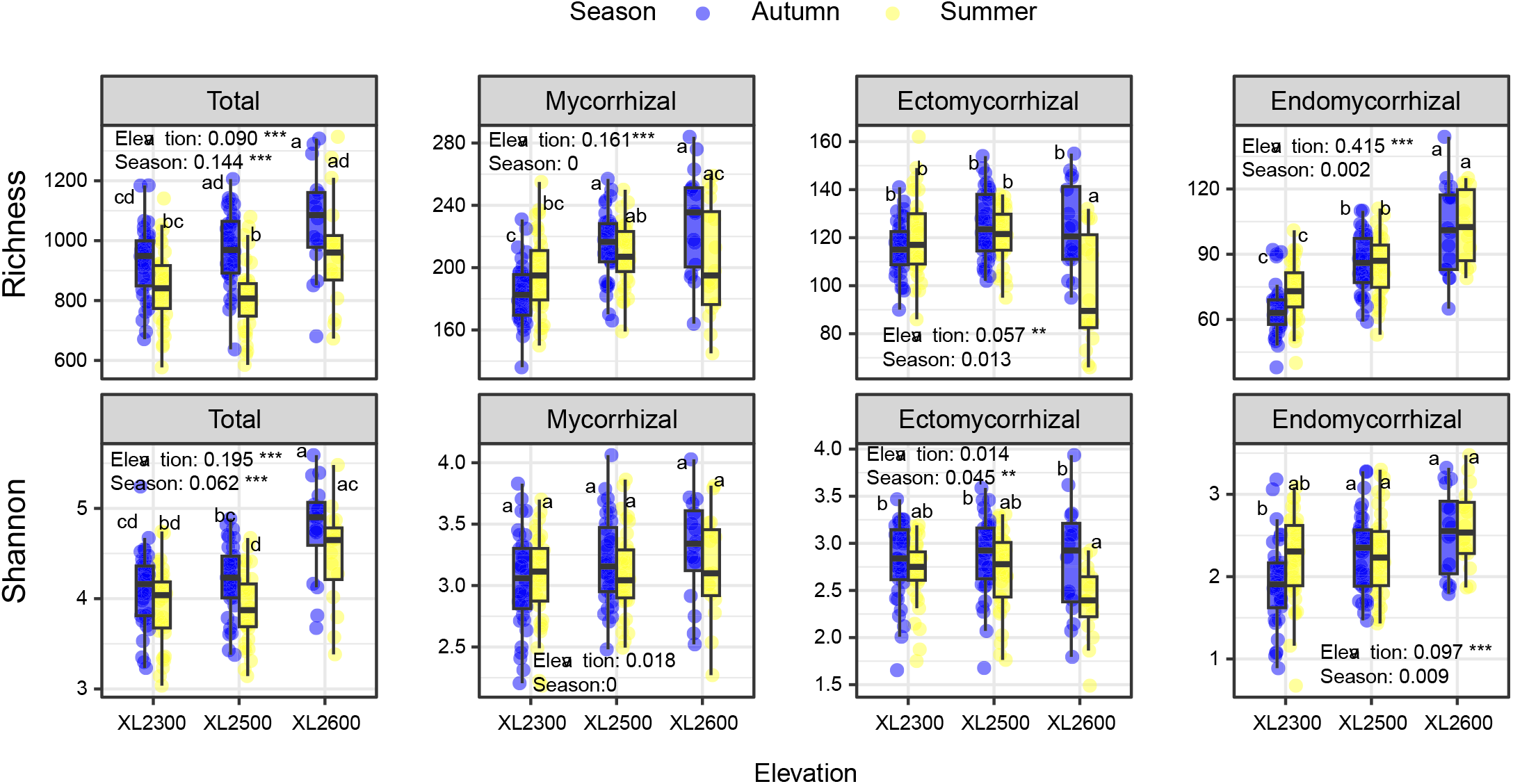
Spatiotemporal distribution of soil fungal richness and diversity during the different elevations and seasons. A two-way analysis of variance was performed to detect the significant effects of elevations and seasons on fungi richness and diversity. The number indicated after the elevation zone and season correspond to *R^2^*, representing the variation in fungal richness and diversity explained by season and zone, *, *P* < 0.05, **, *P* < 0.01, ***, *P* < 0.001. The different superscripts of the box plots were calculated based on the compLetters function in the multcompViewmult package (62) of the Tukey HSD post hoc test.

Non-metric multidimensional scaling(NMDS)and permutation-based multivariate analysis of variance (PERMANOVA) based on the Bray-Curtis distance matrix of all samples indicated that each fungal community composition was significantly separated between elevation (total fungi: *R^2^* = 0.25; total mycorrhizal fungi: *R^2^* = 0.27; ectomycorrhizal fungi: *R^2^* = 0.27; endomycorrhizal fungi: *R^2^* = 0.22. *P* < 0.001), while season only explained a small variance percentage of the fungal community composition (Figure 4A). Although the soil fungal communities in each elevation showed separation in autumn and summer (Fig. S4), these divergences were overthrown by the sampling zones’ impact. In summary, the elevation had a stronger effect on the composition of each fungal community than the seasons.

**FIG 4.**
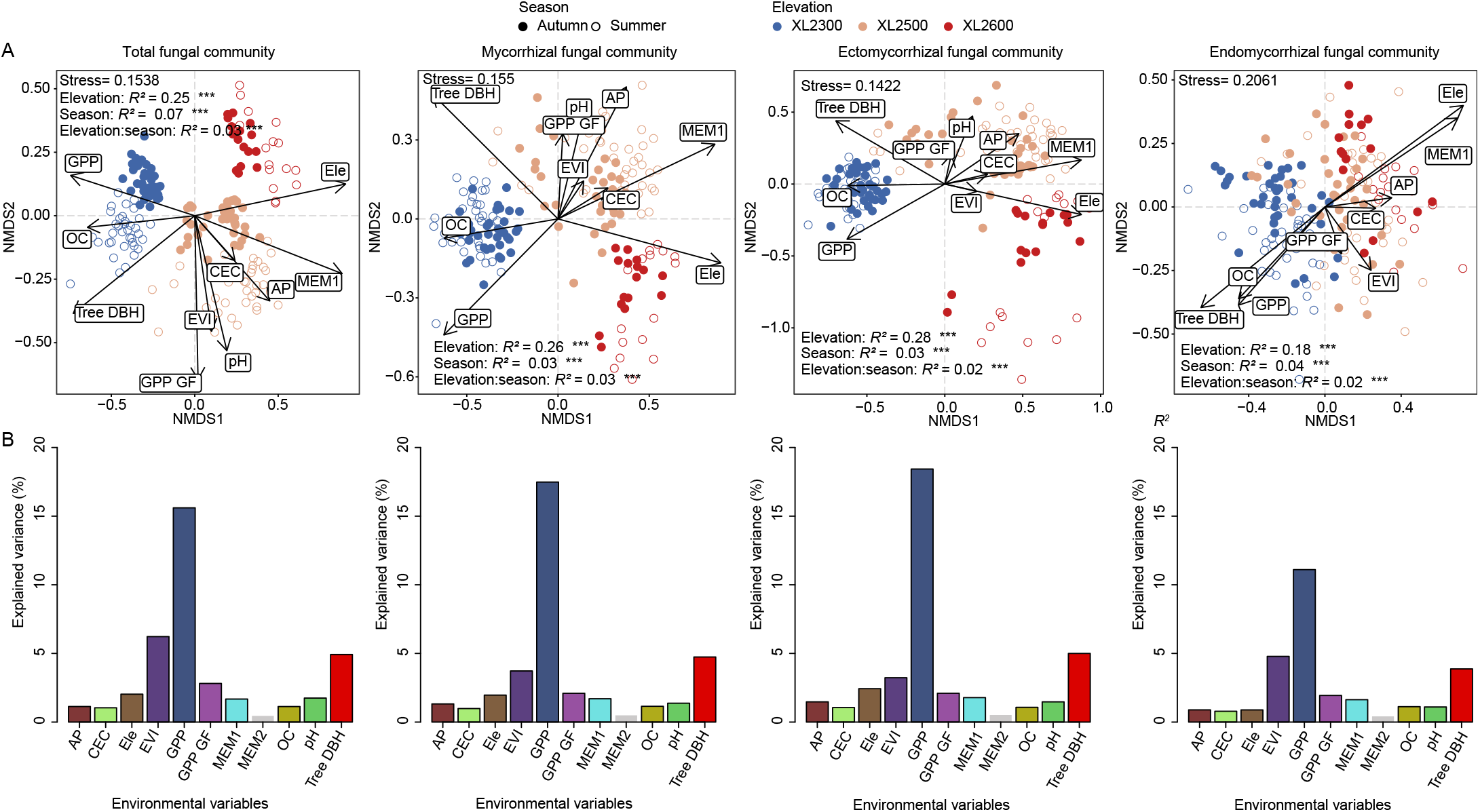
Non-metric multidimensional scaling ordinations (A) based on Bray-Curtis distance matrix and explanation (B) of fungal community composition variation at four levels by environmental variables detected by PERMANOVA. The arrows in the scatter plot indicate the correlation strength and the direction of the maximum increase of the environmental variables with a significant contribution to the community composition variation. The number indicated after the elevation and season correspond to *R^2^*, representing the variation in fungal community composition explained by season and zone, ***, *P* < 0.001. In the bar chart, the bar represents the environmental variables with a significant effect on the fungal community composition variation is edged. AP, soil available phosphorus; CEC, Soil cation exchange capacity; Ele, elevation of sampling point; EVI, enhanced vegetation index; GPP, ground primary productivity; GPP GF, gap-filled of ground primary productivity; MEM, spatial eigenvectors generated from geographic coordinates (latitude and longitude); OC, soil organic carbon; pH, soil acidity and alkalinity; Tree DBH, DBH of the sampled tree.

### Relationship between fungal community composition and geographical and environmental distance

The alteration in fungal community similarity with geographical distance revealed a significant distance-decay relationship (DDR) in fungal community composition. Interestingly, the DDR slope was steeper in the total fungal community, while it was relatively gradual in the functional fungal community, especially the endomycorrhizal fungal community (Figure 5). Moreover, the DDR slope was steeper in summer than in autumn. The similarity of fungal communities decreased with increased environmental distance in autumn, but the variation in summer was lower than in autumn (Fig. S5). The relationship between the fungal community composition similarity and the attenuation of environmental distance was most pronounced in the total fungal community (autumn: *R^2^* = 0.24), and less pronounced in the endomycorrhizal fungal community (autumn: *R^2^* = 0.05). In summary, spatial heterogeneity affected the soil fungal community composition similarity to a greater extent compared to environmental heterogeneity.

**FIG 5.**
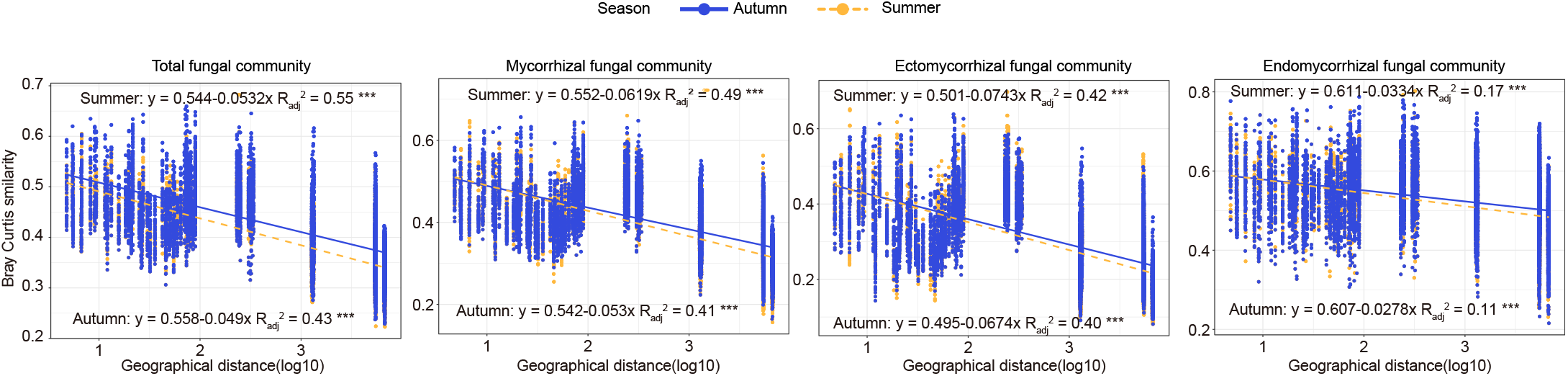
Distance-decay patterns of soil fungal community composition and geographical distance, based on the Bray-Curtis similarity. Different dot colors and line colors and types represent summer and autumn samples. *R^2^*, representing the fitting degree of fungal community similarity and geographical distance, ***, *P* < 0.001.

### Effects of environmental variables on soil fungal diversity and composition

A stepwise multiple linear regression model of fungal richness and diversity with environmental factors was constructed, indicating that environmental variables could only explain 23%, 20%, 11% and 38% of the spatial and temporal variation in richness of the total soil fungi, total mycorrhizal fungi, ectomycorrhizal fungi, and endomycorrhizal fungi. The richness variation in the four fungal communities was synchronously affected by GPP and MEM1, while EVI and pH also contributed significantly. Tree DBH explained a considerable part of the variation of ectomycorrhizal fungal richness (Table 1). More interestingly, tree DBH became more important within the season, while AP also became a major factor influencing richness, and the percentage of variance explained by the model increased (Table S1).

**Table 1.**
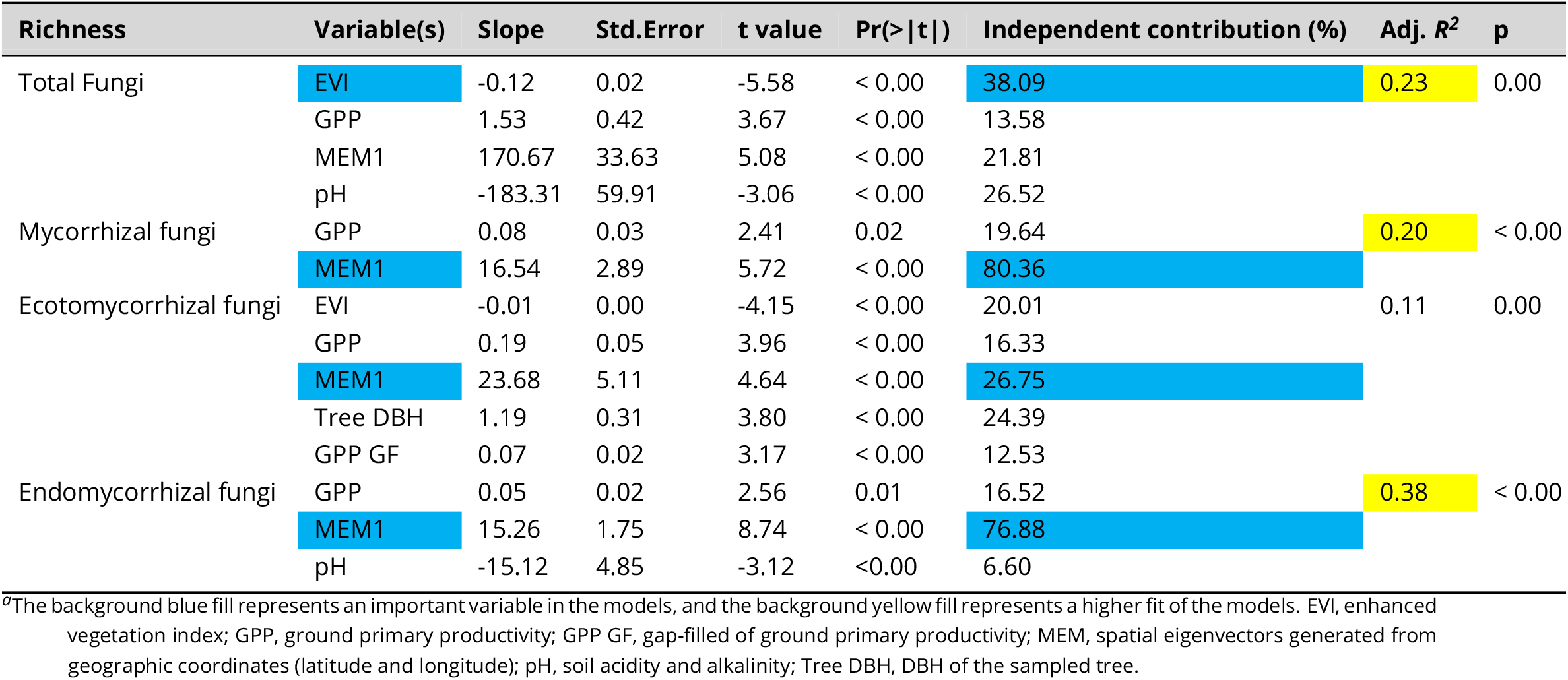
Stepwise multiple regression linear model between soil fungal richness and environmental variables to reveal their contribution to soil fungal richness variation.

Across all zones, the main driver of fungal community composition in the soil was GPP, according to PERMANOVA (total fungi: *R^2^=* 0.15; total mycorrhizal fungi: *R^2^* = 0.17; ectomycorrhizal fungi: *R^2^* = 0.18; endomycorrhizal fungi: *R^2^* = 0.11; *P* <0.01, Figure 4B and Table S2).

Variation partitioning showed that the fungal community composition variance at different levels was explained most consistently by the plant variable (i.e., 34.5%, 32.1%, 31.3%, 33.3% in total, total mycorrhizal, ectomycorrhizal and endomycorrhizal fungal community composition, respectively), followed by the soil variable (i.e., 21.1%, 23.5%, 24.1% and 21.4% in total,total mycorrhizal, ectomycorrhizal and endomycorrhizal fungal community composition, respectively) (Figure 6A). With regards to the fungal richness, the presence of plants variable explains a significant percentage of the variance of each fungal community, especially the total and ectomycorrhizal fungal richness variances (i.e., 8% and 36.8%, in total and ectomycorrhizal fungal richness). Space variable explained the highest variance percentage variance of total mycorrhizal fungal richness (28.7%), while elevation explained the highest variance percentage of endomycorrhizal fungal richness (31.1%) (Figure 6B).

**FIG 6.**
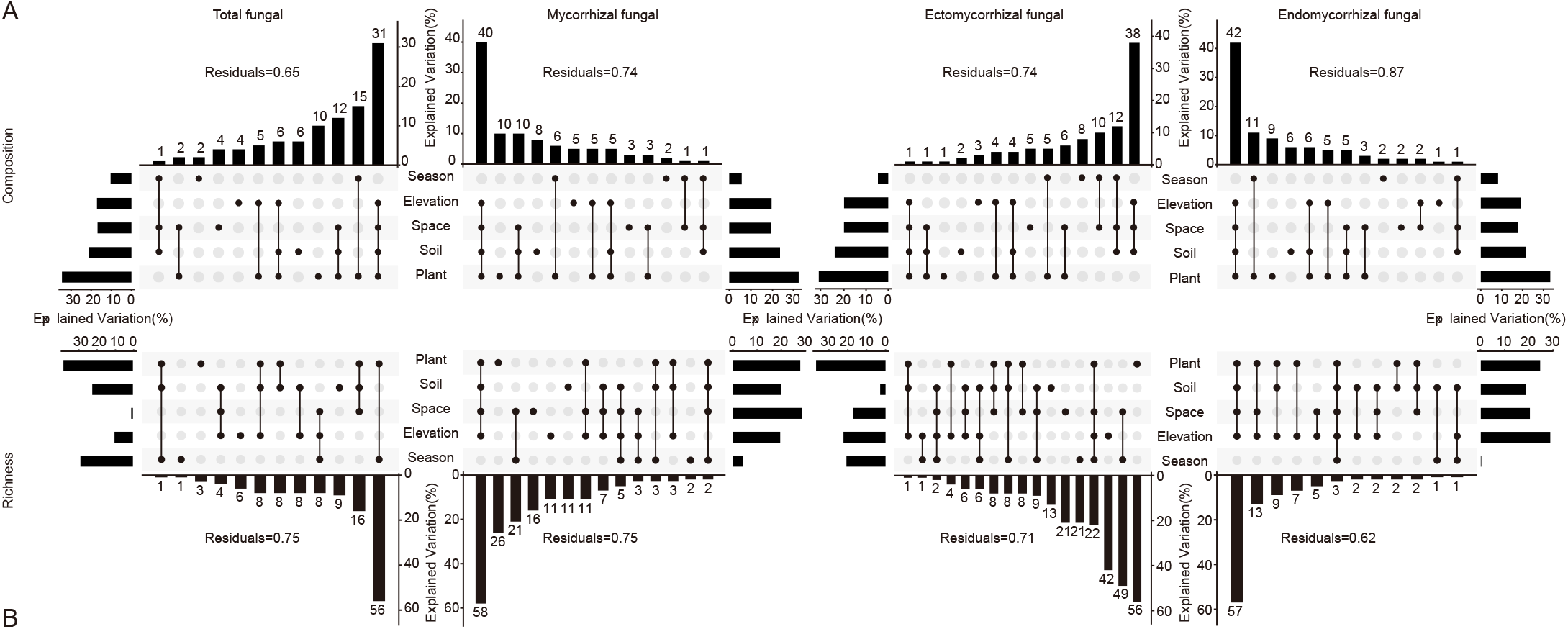
Upset plot of the hierarchical variation partitioning results, showing the individual and shared contributions of season, elevation, space, soil, and plant on soil fungal richness (B) and community composition (A), respectively. The dot matrix and histogram present the values for shared and exclusive contributions, and the horizontal histogram presents the percentage of individual effects toward the total explained variation. Residuals represent the percentage unexplained by these variables.

In summary, the environmental variables could only explain a small percentage of the variance in soil fungal diversity and community composition. Different types of environmental variables exhibited different percentages of the fungal richness variance explained, while the causes for the variance of different fungal community compositions were relatively consistent.

### Ecological process of fungal community assembly

We explored the distribution of *β*NTI in four fungal communities to infer the ecological processes of soil fungal communities assembly. We observed sign ificant differences between the four fungal communities (Figure 7A). Both deterministic and stochastic processes considerably influenced the assembly of the total fungal community. They showed seasonal differences (deterministic processes: 47% in summer and 62% in autumn, stochastic processes: 53% in summer and 38% in autumn). On the other hand, the aggregate of the other three fungal community levels was more affected by a random process and did not exhibit seasonal differences (mycorrhizal fungi: 89% in summer and 86% in autumn, ectomycorrhizal fungi: 90% in summer and autumn 92%; endomycorrhizal fungi: 71% in summer and 73% in autumn).

**FIG 7.**
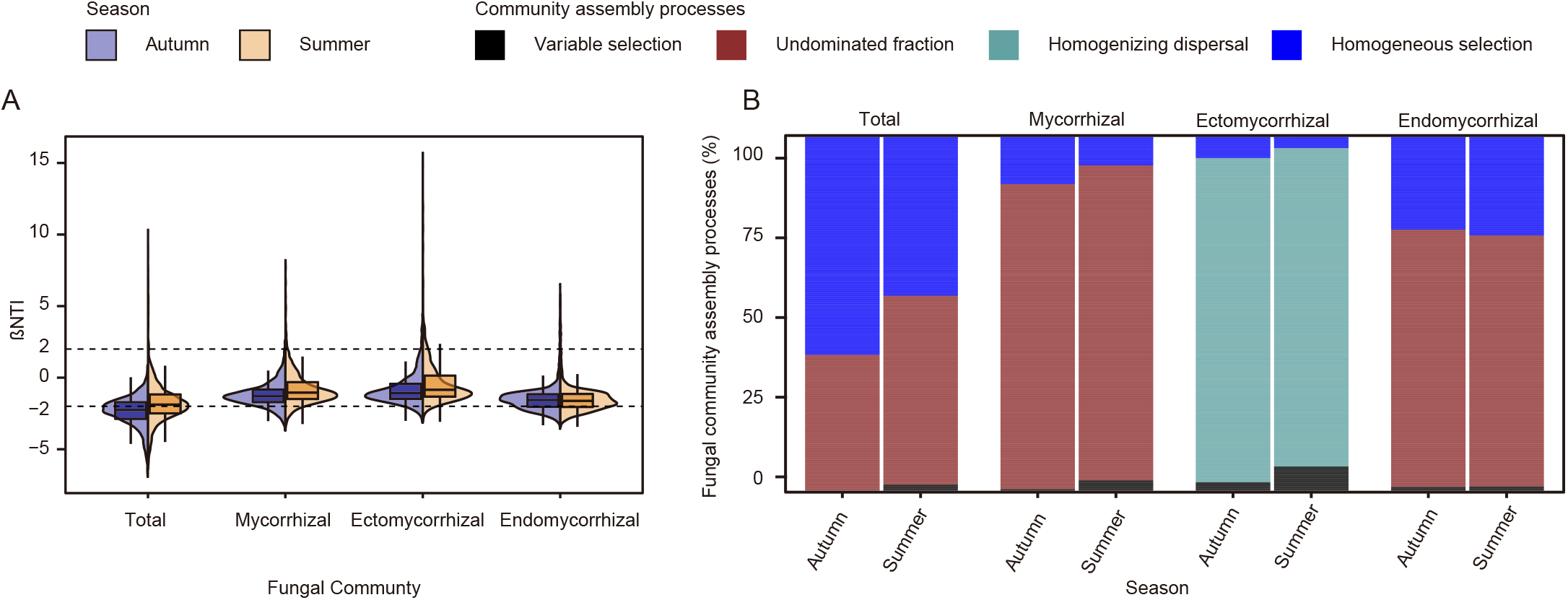
Distribution patterns of *β*NTI in soil fungal communities at four levels (A) and ecological processes driving soil fungal community assembly (B).

With regards to the more detailed ecological processes of the fungal community assembly, they differed significantly at each level: the strong homogeneous selection (50%) and weak variable selection (1%) in the deterministic process and the undominated fraction (49%) in the stochastic process determined the total fungal community assembly. The strong undominated fraction (89%) in the stochastic process and the relatively weak homogeneous selection (10%) and variable selection (1%) in the deterministic process determined the total mycorrhizal fungal community assembly. The strong homogenizing dispersal (91%) in the stochastic process and a weak homogeneous selection (5%) and variable selection (4%) in the deterministic process determined the ectomycorrhizal fungal community assembly. Finally, the strong undominated fraction (77%) in stochastic process and the strong homogeneous selection (21%) and weak variable selection (1%) in the deterministic process determined the endomycorrhizal fungal community assembly. In general, except for the fact that the assembly of ectomycorrhizal fungal communities was strongly affected by homogenizing dispersal, the other three fungal communities assemblies were affected by homogeneous selection, undominated fraction and variable selection, but the proportions were completely different (Figure 7B).

## DISCUSSION

The mechanisms underlying the spatial and temporal distribution mechanism of soil microorganisms have been a long-standing but controversial issue. This work has explored the diversity patterns and assembly mechanisms of soil fungal communities on two important spatial and temporal gradients (elevation and season) at a regional scale. This work focused on *P. davidiana* forests. More importantly, this work investigated in detail the different levels of fungal communities in soil. The results showed that the elevation was the dominant factor affecting the spatial and temporal variation of the soil fungal community, and it had variable effects on the soil fungal community at different levels. However, neither the elevation nor the season led to a large-scale variation of soil fungal communities. Further, we found that plant variables mainly explained the spatial and temporal variation of the soil fungal community. The fungal community composition showed a significant distance decay pattern. In addition, we found that the stochastic processes were more dominant compared to the deterministic process (i.e., variable selection) in shaping soil fungal community composition in the *P. davidiana* forest. The relative contribution of ecological processes varied among the different soil fungal communities.

### Spatiotemporal dynamics of the soil fungal community on Xinglong Mountain

The soil fungal alpha diversity variation among elevations was greater than that among seasons, especially the richness of endomycorrhizal fungi, exhibiting a significant increase with increasing elevation. It is generally suggested and reported that fungal richness decreases with increasing elevation (63, 32, 61), and it has also been reported that fungal richness does not increase with elevation (33). Therefore, the explanation of the elevation model of endomycorrhizal fungal richness is also diverse. It is generally believed that endomycorrhizal fungi are more likely to occur at the seedling stage of mycorrhizal plants (64). We speculated that with the increase of elevation, the decreasing maturity of the populus forests leads to an increase in the richness of endomycorrhizal fungi. In contrast, the variation of ectomycorrhizal fungal richness between elevation was significant but relatively small. This may be because the three elevation zones are poplar forests with the same host tree species. The host tree species usually have a strong effect on ectomycorrhizal fungi variation (4, 44, 4).

Compared with the significant variation between elevation, the soil fungal richness variation between seasons was only significant for the total fungal richness. This was mainly attributed to the fact that the total fungal richness in autumn was significantly higher than that in summer, similar to previous reports (36, 37). In contrast, the richness of total mycorrhizal fungi, ectomycorrhizal fungi, and endomycorrhizal fungi did not show a significant seasonal pattern (Figure 3). Our results indicated that the symbiotic fungi richness is not affected significantly by seasonal changes. In fact, studies have shown that the ectomycorrhizal fungal richness increases significantly in April and remains fairly stable until October (65). Other studies spanning a longer period than comparing mycorrhizal fungal richness in autumn and summer have reported seasonal variations (38, 39, 40), but this does not conflict with our findings. As for the seasonal stability of mycorrhizal fungal richness identified in this study, we suggest that it might be a result of small fluctuations of soil physical and chemical variables between seasons. The total fungal community is usually affected by physical and chemical properties fluctuations, such as pH and AP (66, 67), but we assume that these soil variables do not significantly change with seasons, so we cannot provide specific conclusions.

In studying the spatial and temporal variation of soil fungal communities at large spatial scales, the spatial variation is usually greater than the variation between seasons (68, 69). Although our research was carried out on a regional scale, the results still showed that the soil fungal community composition temporal and spatial variation in a *P. davidiana* forest was mainly dominated by elevation. Although the season also significantly shaped the soil fungi community composition, it had a less significant effect on soil fungi than the elevation.

### Effects of geographical distance on soil fungal community composition

Microbiological studies at different spatial scales have shown that microbial communities are usually affected by the geographical distance on a larger scale and by environmental aspects on a shorter, regional scale (70, 71). However, our results showed that, at the regional scale, the spatial heterogeneity of soil fungal community composition could also be significantly explained by the distance decay relationship (DDR) (Figure 5). The DDR indicated that the variation in the composition of the four fungal communities was significantly correlated with geographical distance. The DDR slope of endomycorrhizal fungal community composition was the lowest, but this did not correspond to a lower turnover rate of endomycorrhizal fungi associated with *P. davidiana* with regard to geographical distance. This might be due to the high migration of fungal species weakening the DDR by homogenizing the community (72). Thus the low DDR slope observed for the endomycorrhizal fungal community might be due to its high migration ability. The null model in our study indicated that the endomycorrhizal fungal community migrates through homogeneous selection and thus weakens DDR (Figure 7). In addition, the four fungal communities had a weaker DDR in autumn was weaker compared to the summer, indicating that the turnover of soil fungi in autumn was lower than that in summer. The reason underlying the low slope of DDR in autumn should be similar to that explaining the low slope of DRR of endomycorrhizal fungi: homogeneous selection of fungal communities with high relative importance in autumn led to high migration of fungi and weakened the DDR.

### Assembly of soil fungal community

Variation partitioning helps us understand the effects of environmental variables on the fungal richness and fungal community composition. Environmental variables only explained a small part of soil fungal communities’ spatial and temporal variation. Multiple sets of environmental variables explained this portion of the variation, and the portion explained by pure elevation or season was even less. Plant variables greatly contribute to the shaping of fungal community composition. Although our results show that plant and other environmental variables explain a part of the variation of fungal richness and community composition, a large part of the variation cannot be explained. This unexplained variation is generally caused by noise in the ecological process of microbial community assembly (52, 53, 73, 74). We evaluated the ecological processes of fungal community assembly using null models. βNTI and Raup-Crick were used to determine the assembly process of each fungal community. Notably, except for the ectomycorrhizal fungal community, the assembly of the other three fungal communities was controlled by undominated process and homogeneous selection. Homogeneous selection accounted for a large proportion of the ecological process, especially in the total and endomycorrhizal fungal communities, while the ectomycorrhizal fungal community assembly was mainly driven by homogenizing dispersal. Homogeneous selection refers to the process by which the environment limits the microbial populations’ differentiation and is an ecological factor that alters community structure in a homogenous state, resulting in similar community structures with deterministic variables. Such factors include biotic and abiotic conditions (51). The homogeneous selection was dominant in the total and endomycorrhizal fungal community assembly process, indicating that in the range of environmental heterogeneity in this study, it decreased the fungal community differences and also homogenized the fungal community and endomycorrhizal community, resulting in a high similarity of soil fungal communities (51). There is always a process of variable selection (heterogeneous selection) in the ecological process of four fungal community assembly, but the proportion is very small, which may be the reason for the diversity of fungal community in different environments, similar to the small variation of fungal community composition explained by environmental variables in the results of variance partitioning (Figure 5). Dispersal limitation is considered to be due to certain limitations in the migration of organisms, such as spatial distance and environmental filtering. Unlike other reports on the dominant role of dispersal limitation and variable selection in forest soil fungal communities (55, 51), the dispersal of the four fungal communities in this study was not limited. In particular, the ectomycorrhizal fungal communities assembly was very different from the other three fungal communities, mainly dominated by homogeneous dispersal, indicating the strong dispersion of ectomycorrhizal fungi in *P. davidiana* soil. The environmental heterogeneity in this study was at the limit of ectomycorrhizal fungal adaptation range, but due to its strong dispersal, it has not formed a heterogeneous community structure (51). The undominated fraction (including weak selection, weak dispersal, diversification and drift) is caused by the species’ random birth, death, and reproduction, which is not related to niche preference. The undominated fraction is important in assembling communities other than ectomycorrhizal fungal communities. In combination with the dominant role of homogeneous selection in the assembly of these three fungal communities, we conclude that homogeneous selection and ecological drift are more important than the niche-related environmental selection at the regional scale of this study (75).

### Conclusions

We assessed the spatial and temporal distribution of soil fungi in the Xinglong Mountain forests dominated by *P. davidiana*. The richness of the different fungal communities exhibited different spatial or seasonal patterns, but the composition of these communities was mostly affected by the spatial patterns compared to the seasons. This may be because the environmental heterogeneity caused by space was greater than the niche difference between seasons. The spatial and temporal distribution patterns of community composition of different soil fungi types could be explained by environmental variables, especially plant variables. At the same time, different environmental variables explained the spatial and temporal patterns of the richness of different types of fungi. All four fungal communities showed a significant DDR, indicating a high turnover rate. The assembly of total mycorrhizal and endomycorrhizal fungal communities showed a higher proportion of undominated fraction (including weak selection, weak dispersal, diversification and drift), while the assembly of total fungal communities was controlled by homogeneous selection and undominated fraction, and the assembly of ectomycorrhizal fungal communities was dominated by homogeneous dispersal. Variable selection (heterogeneity selection) played a minor role in the four fungal community assembly in this study, and dispersal limitation did not exist. At the regional scale, environmental heterogeneity did not lead to a dramatic variation of fungal communities. Still, environmental heterogeneity, especially plant variables, was a reasonable explanation for fungi’s spatial and temporal variation. The study evaluated elevation differences, but no clear elevation patterns were observed, which may be caused by small elevation variations or insufficient gradient. This study highlighted the changing patterns and ecological processes of forest soil fungal communities dominated by dual-mycorrhizal plants, especially symbiotic mycorrhizal fungal communities, and improved our understanding of the integrity and diversity of soil fungal communities. Therefore, future studies should investigate soil fungal communities in a comprehensive and differentiated manner to provide more valuable results.

## MATERIALS AND METHODS

### Sampling, soil physicochemical properties, tree characteristics, and climate data

The soil samples were collected from the Xinglong Mountain National Nature Reserve, located about 45 km southeast of Lanzhou City (103°50’-104°10’E, 35°38^/^-35°58’N), with an elevation of 1800-3670 m. The region has a temperate semi-humid and semi-arid climate type (76). In the suitable elevation range of *Populus davidiana* distribution in the Xinglong Mountain area, three sampling zones were selected, corresponding to a low elevation (XL2300)(104°3’59”E, 35°47’56”N, 2,317 m to 2,344 m above sea level), a middle elevation (XL2500)(104°3’15”E, 35°45’5”N, 2,529 m to 2,532 m above sea level) and a high elevation (XL2600) (104°2’49”E, 35°44’28”N, 2,613 m to 2,615 m above sea level), respectively. The sampling was conducted in the summer (June 2020) and autumn (September 2020). In XL2300 and XL2500 zones, three plots were set in each zone, and three independent trees were selected for sampling from each plot (more than 5 meters apart). In XL2600 zones, only one plot containing four independent sample trees was set. The distance between the sampled trees in each zone was at least 5 m. The DBH of each sample tree was measured, and photographed its growth to obtain its characteristic information (Figure 1). Soil cores of 20 cm depth were drilled from the topsoil in four directions (east, south, west, and north), 1.5 m to 2 m away from the trunk of the tree. Each soil sample was passed through a 2 mm soil sieve, and then it was stored in 50 ml and 2 ml sterile centrifuge tubes. The samples were transported back to the laboratory using dry ice and stored at −80°C for molecular analysis. The remaining soil of each sample was air-dried for analysis of the physical and chemical properties.

The soil samples’ physical and chemical properties were only measured in the summer because the physical and chemical properties will not change greatly in a short period (77). In addition, the four samples of each tree were combined into two samples (southeast and northwest), meaning that soil samples from the east and south shared physical and chemical property information, as did samples from the west and north. The soil’s physical and chemical properties were determined by Baisheng Biotechnology Co., Ltd., Xilin Gol League, Inner Mongolia, according to China’s agricultural and forestry industry standards. Specifically, soil total nitrogen (TN) was measured using the Kjeldahl method, soil organic carbon (OC) was measured using the potassium dichromate volumetric method, soil available phosphorus (AP) was measured using the molybdenum antimony anti-colorimetric method, soil cation exchange capacity (CEC) was measured using the ammonium acetate exchange Kjeldahl method, and soil pH was measured using the acidity meter method.

Climate data were available on the freely accessible website database Worldclim (https://www.worldclim.org/data/index.html). The GPP and EVI data of each zone were extracted and used as a proxy for the zone’s total primary productivity and above-ground net productivity using the MOD17A2H product with a spatial resolution of 500 m 500 m and an 8-day temporal resolution and the MOD13Q1 product with a spatial resolution of 250 m 250 m and a 16-day temporal resolution provided by the MODIS-Tools package (78). The *dbmem* function in the adespatial package (79) was used to construct a distance-based Moran’s eigenvector map (dbMEM) from the latitude and longitude coordinates of sampling points.

### Molecular analyses

Total soil DNA extraction from 50 mg of soil samples was performed using the Qiagen DNeasy PowerSoil DNA Isolation Kit (Qiagen, Germany) following the manufacturer’s instructions. Each sample was extracted in triplicate, and the total DNA quality and quantity were evaluated using a NanoDrop ONE spectrophotometer (Thermo Scientific, USA) and pooled for subsequent analyses. We used three primer pairs to amplify different regions of the soil microbial DNA. The fungal internal transcribed spacer region 1 (ITS1)-targeting primer pairs were ITS1F (5’-CTTGGTCATTTAGAGGAAGTAA-3’)/ ITS12 (5’-GCTGCGTTCTTCATCGATGC-3’) (80); the primers ITS86 F (5’-GTGAATCATCGAATCTTTGAA-3’)/ ITS4R (5’-TCCTCC GCTTATTGATATGC-3’) (81) were used to target the internal transcribed spacer region 2 (ITS2), and primer pairs AMV4.5NF (5’-AAGCTCGTAGTTGAATTTCG-3’)/ AMDGR (5’-CCCACTATCCCTATTAATCAT-3’) were used to amplify a fragment of the arbuscular mycorrhizal fungi (AMF) 18S rRNA gene region (82). The 30 μl PCR reaction system contained 15 μl of Phusion high-fidelity PCR Master Mix (New England Biolabs), 0.2 μM forward and reverse primers, and 10 ng of template DNA. Amplification was performed as follows: 1 min initial denaturation at 98°C, 30 cycles of 10 s at 98°C, 30 s at 50 °C, and 30s at 72°C, with a final 5 min elongation at 72°C. Following the manufacturer’s instructions, libraries were generated using the Illumina TruSeq DNA PCR-Free Library Preparation Kit (Illumina, USA), and index codes were added. The Qubit 2.0 Fluorometer from Thermo Scientific and the Agilent Bioanalyzer 2100 system was used to evaluate the library’s quality. All samples were pooled into equimolar concentrations before sequencing with the paired-end protocol on the Illumina NovaSeq platform by Novogene Biotech Co., Ltd (Tianjin, China).

### Sequencing Statistics

Raw sequences were split into groups based on their barcodes. The paired-end raw sequences were processed in the QIIME2 platform (83). The Cutadapt plugin was used for primers removal from paired-end sequences, and the DADA2 denoise-paired plugin was used for sequence quality control of paired-end reads, and amplicon sequence variants (ASVs) clustering with 100% similarity was obtained. Operational taxonomic units (OTUs) were obtained by clustering the ASVs based on a 97% identity threshold of the sequences using the q2-vsearch plugin. OTUs present in only one sample was removed. The Qiime feature-classifier classifier-sklearn pipeline was used to classify OTUs to identify their taxonomic ranks. Reference sequences for training the sciKit-learn naive_bayes classifier were obtained from UNITE version 4 (84) and MaarjAM databases (85). The FUNGUild v1.1 script (86) was used to predict the OTUs function, and different types of total mycorrhizal fungi OTU were screened based on the results. Based on FUNGuild prediction results, all OTUs in this study were divided into four fungal communities: the total fungal community, the total mycorrhizal fungal community, the ectomycorrhizal fungal community, and the endomycorrhizal fungal community. It should be noted that the total mycorrhizal fungal community OTUs were defined as OTUs that were predicted to be from mycorrhizal fungi and contained all mycorrhizal fungal types. The ectomycorrhizal fungal community OTUs were defined as the OTUs that were predicted to be from ectomycorrhizal fungi, and these were preferentially considered ectomycorrhizal fungi. The endomycorrhizal fungal community OTUs were defined as the OTUs from the total mycorrhizal fungal community other than the ectomycorrhizal fungi.

### Statistical analyses

The *chart.Correlation* function in the PerformanceAnalytics package (87) was used to assess the normal distribution of environmental variables and the pairwise correlation between variables. The soil physical and chemical properties variable TN and the climate variables Avetmax and Aveorec were removed because these three variables were co-linear with other environmental variables (r > 0.7). Since the environmental variables did not follow a normal distribution (Fig. S1), we used non-parametric methods to evaluate the relative importance of the elevation and season to environmental variables. The *kruskal_test* function in the rstatix package (88) was used to evaluate the effects of the elevation on four physical and chemical properties of soils and the DBH of host trees. The *scheirerRayHare* function in the rcompanion package (89) was used to evaluate the effects of elevation and season on EVI, GPP, and GPP GF.

The rarefaction and alpha diversity calculation of the fungal community datasets was performed using the vegan packag *rrarefy, estimateR*, and diversity functions (90). A two-way ANOVA was used to evaluate the effect of different seasons and altitudinal regions on alpha diversity, and pairwise comparisons were performed using Tukey’s multiple comparison method. Based on the two-way ANOVA results, a linear regression model was fitted to elevation and fungal abundance to accurately assess the importance of elevation on fungal richness. A stepwise multiple linear regression was used to explore the multivariate explanation of the pattern of fungal richness variation pattern, and each variable’s independent contribution was calculated using the *hier.part* function in the Hier.part package (91).

The *vegdist* function from the vegan package (90) was used to calculate the Bray-Curtis distance matrix for the community datasets of total soil fungi, total mycorrhizal fungi, ectomycorrhizal fungi, and endomycorrhizal fungi. In this study, fungal communities’ dissimilarities were ordinated using the non-metric multidimensional scaling (NMDS) method based on the Bray-Curtis distance matrix. To determine the contribution of the two experimental factors (different seasons and different elevations) to the soil fungal community structure in this study, we analyzed the Bray-Curtis distance matrix between pairs of samples with a permutation-based test using a PERMANOVA model of the *adonis* function. To determine the importance of geographic distance on the fungal community similarity, linear models of the geographic and environment distance matrix of sampling points and the Bray-Curtis similarity matrix of the fungal community were fitted. The geographic distance matrix between sampling points was obtained by calculating the latitude and longitude coordinates data of sampling points using the *distm* function in the geosphere package (92). The environmental distance matrix was the Euclidean distance between zones based on measured environmental variables.

To evaluate the effects of environmental variables on soil fungal community composition, we first used the *envfit* function in the vegan package (90) to fit the environmental variables with the NMDS results. Then we used PERMEANOVA to quantify the effects of various variables on soil fungal community composition. In addition, to quantify the relative importance of different environmental variables on the variation of soil fungal richness and community composition, the *rdacca.hp* function from the rdacca.hp package was used to perform hierarchical, and variation partitioning on the total variation of soil fungal richness and community composition explained by environmental variables (93). The environmental variables were divided into three types: soil (OC, AP, pH, CEC), plant (EVI, GPP, GPP GF, tree DBH), and space (MEM1, MEM2). Together with elevation and season, they were used for variation partitioning and total variation hierarchy.

The *pNST* function in the NST package was used to calculate the *β*-nearest taxon index (*β*NTI) between paired samples and the Bray-Curtis-based Raup-Crick metric (RCbray) (94), and the community assembly process was inferred using the previously developed null model (95, 96, 97, 98) specifically if the observed *β*MNTD value does not deviate significantly from the null *β*MNTD distribution (|*β*NTI| < 2), it indicates that the phylogenetic composition differences in the observed community are due to uncertain processes (including diffusion limitation, homogenization diffusion). If the *β*NTI value < −2, the observed community phylogenetic development is significantly lower than the expected phylogenetic replacement (that is, the community assembly is driven by homogeneous selection). If *β*NTI > 2, it indicates a significantly higher than the expected system replacement (that is, the community assembly is driven by variable selection). At the same time, based on the method first proposed by Stegen and modified by Stegen and Dini Andreote et al., we performed a more detailed assessment of the community assembly process when |*β*NTI| < 2: when |*β*NTI| < 2 and RCbray > 0.95, the community assembly between samples will be considered as dispersal limitation; when |*β*NTI| < 2 and RCbray < −0.95 between paired samples, the community assembly between samples will be considered as homogenizing dispersal; when |*β*NTI| < 2 and RCbray < 0.95, the community assembly between samples will be considered as an undominated fraction (including weak selection, weak dispersal, diversification and drift).

### Availability of data and materials

Raw sequences were deposited in the Sequence Read Archive under Bioproject PRJNA852440. All supplemental figures and tables that appear in the text were organized in a collection document SUPPLEMENTAL FILE1. The read count OTU table and the representative sequence of each OTU were provided in SUPPLEMENTAL FILE2. The corresponding metadata was provided in SUPPLEMENTAL FILE3.

## SUPPLEMENTAL MATERIAL

Supplemental material is available online only.

**SUPPLEMENTAL FILE1**, PDF file, 1.28 MB.

**SUPPLEMENTAL FILE2**, XLSX file, 6.58 MB.

**SUPPLEMENTAL FILE3**, XLSX file, 31 KB.

## ACKNOWLEDGMENTS

We would like to acknowledge Dr. Zhao Heng for his advice in data analysis, and Dr. Wu Fang for her help in manuscript writing. This study was supported by the National Natural Science Foundation of China (Project No. U1802231), the China Postdoctoral Science Foundation (2019M660508 to QZ) and the Beijing Tree Molecular Design Breeding Advanced Innovation Center and the ARBRE Excellence Laboratory (ANR-11-LABX-000201).

